# Histone variant H2A.Z mutant suppresses the senescence-associated secretory phenotype

**DOI:** 10.64898/2026.02.16.706244

**Authors:** Zong Ming Chua, Hiroshi Tanaka, Adrianna Abele, Adarsh Rajesh, Marcos Garcia Teneche, Xue Lei, Karl Miller, Andrew Davis, Carolina Cano Macip, Laurence Haddadin, Nirmalya Dasgupta, Peter D. Adams

## Abstract

Cellular senescence features a durable cell-cycle arrest and a pro-inflammatory senescence-associated secretory phenotype (SASP) driven in part by chromatin remodeling. The histone variant H2A.Z plays an essential role in regulating gene expression through modulating nucleosome dynamics and is known to regulate the expression of cell cycle genes during the early stages of cellular senescence. However, how the intrinsic stability of H2A.Z-containing nucleosomes plays a role in the establishment of the senescent phenotype remains unexplored. To investigate this, we employed H2A.Z R80C, a H2A.Z mutant that destabilizes nucleosomes by disrupting histone-DNA interactions. We observed that expression of H2A.Z R80C causes suppression of SASP in senescent primary human fibroblasts, without affecting expression of cell cycle genes. H2A.Z knockdown did not suppress the SASP, demonstrating that SASP suppression is likely due to altered stability of H2A.Z-containing nucleosomes rather than loss of H2A.Z function. Mechanistically, SASP suppression is linked to decreased H3K27ac at SASP gene loci. These findings offer a novel avenue for understanding and manipulating the SASP during aging and other senescence-related pathologies.

## INTRODUCTION

Cellular senescence, a state of irreversible cell cycle arrest triggered by both extrinsic and intrinsic stimuli, is a complex biological process marked by significant alterations in chromatin structure, epigenetic modifications, metabolism and gene expression^1-6^. Senescent cells remain viable after stable cell cycle arrest, and their burden is known to increase with age in many vertebrates^7^. Senescent cells produce a specific pro-inflammatory senescence-associated secretory phenotype (SASP) which consists of proinflammatory cytokines such as IL-6 and IL-1, chemokines such as IL-8, and extracellular proteases such as MMP-1. This SASP can improve tissue repair and regeneration, anti-tumor immunity, and cellular plasticity^8-13^. However, sustained and chronic expression of the SASP, especially in older organisms, can result in the spread of senescence and impaired tissue function through paracrine signaling^14^. Chronic SASP signaling can also lead to a pro-inflammatory and immunosuppressive microenvironment that drives tumorigenesis^15^. Accordingly, removal of senescent cells has been shown to improve tissue function and promote healthy aging^16 17^

Induction of cellular senescence and the SASP takes place concurrently with changes in nucleosome composition and localization of histone variants, which in turn regulate key age-related and senescence phenotypes^18-23^. Among the histone variants that play a role in regulating the senescent phenotype is H2A.Z, a variant of core histone H2A, which has been shown to accumulate with age and is linked to age-related cognitive decline in mice^24^. H2A.Z has a roughly 60% degree of sequence similarity with canonical H2A, with a higher proportion of residues in the N- and C-terminal tail domain showing differences compared to the more conserved core regions^25^.

In some contexts, H2A.Z has a pro-proliferative, anti-senescence role. The p53-dependent localization of the histone variant H2A.Z by the p400 complex represses basal p21 transcription in proliferating cells, and upon DNA damage, H2A.Z is evicted to permit Tip60 recruitment and p21-mediated proliferation arrest^26^. Additionally, H2A.Z supports the expression of proliferation-associated genes such as Myc and Ki-67, further reinforcing its role in cell cycle progression and senescence avoidance^27^. In pancreatic ductal adenocarcinoma (PDAC), overexpression of H2A.Z enhances tumor growth and confers chemoresistance by suppressing senescence, again highlighting a pro-proliferative function^28^. On the other hand, H2A.Z is expressed throughout the cell cycle and in non-proliferating cells, unlike canonical H2A which is predominantly expressed during S phase^29^. Accordingly, it has been shown that the growth-arrested senescent phenotype is associated with the site-specific deposition of H2A.Z, suggesting a role for H2A.Z in non-proliferating senescent cells.

The histone variant H2A.Z directly impacts the structure and the stability of assembled nucleosomes, altering the activity of chromatin remodelers and the accessibility of the underlying DNA, thereby affecting diverse DNA-related processes such as repair, recombination, and transcription^30-33^. H2A.Z is localized at the transcription start site (TSS) of actively transcribed genes, and its presence plays a role in regulating the activity of poised genes which are marked by both activating and repressive histone marks^34,35^. Homotypic nucleosomes containing pairs of H2A.Z histones show decreased nucleosomal stability, while H2A.Z containing nucleosomes show altered dynamics, mobility, and increased accessibility of the local chromatin to transcriptional machinery^36-39^.

Given both the importance of cellular senescence and the accompanying SASP in aging, and the profound changes in transcriptome that take place during the onset of cellular senescence and the proposed role of H2A.Z in control of gene expression through nucleosome stability, we sought to investigate how modulating the stability of H2A.Z containing nucleosomes would impact the SASP, potentially providing new avenues to ameliorate age-associated disease phenotypes.

To understand how the stability of H2A.Z-containing nucleosomes plays a role in the induction of cellular senescence, we examined the effects of a specific histone mutant H2A.Z R80C in senescent cells. H2A.Z R80C was previously identified as a histone mutant recurrently found in some cancers (a so-called oncohistone mutation) and has been shown to decrease the stability of the core nucleosomal complex^40,41^. Modeling of the H2A.Z R80C mutant shows that removal of the arginine 80 residue reduces the affinity between the H2A.Z histone and the adjacent DNA strand, as the arginine interacts favorably with the negatively charged DNA backbone while also extending into the minor groove of the DNA strand to stabilize the nucleosome^41^. We used H2A.Z R80C as a tool to probe the relationship between nucleosome stability and phenotypes of cell senescence.

## RESULTS

### H2A.Z R80C alters nucleosome stability

To investigate the relationship between H2A.Z stability and phenotypes of senescence, we employed the H2A.Z R80C mutant that subtly destabilizes nucleosomes^40^. The H2A.Z R80C substitution mutation replaces an arginine residue with cysteine, which is modeled to destabilize the nucleosome due to the loss of the favorable arginine-DNA interaction in an assembled nucleosome (Figure 1A). We carried out biochemical characterization of lentivirus-mediated ectopically expressed H2A.Z wild type (WT) or mutant H2A.Z R80C, both with an N-terminal HA-tag or GFP-tag, in IMR-90 primary human fibroblasts. We confirmed that ectopic H2A.Z WT and R80C were expressed at comparable levels, based on western blotting and quantification of the immunofluorescent signals (Figure 1B, 1C). More than 99% of cells showed clear ectopic expression of H2A.Z WT or R80C.

**Figure 1.**
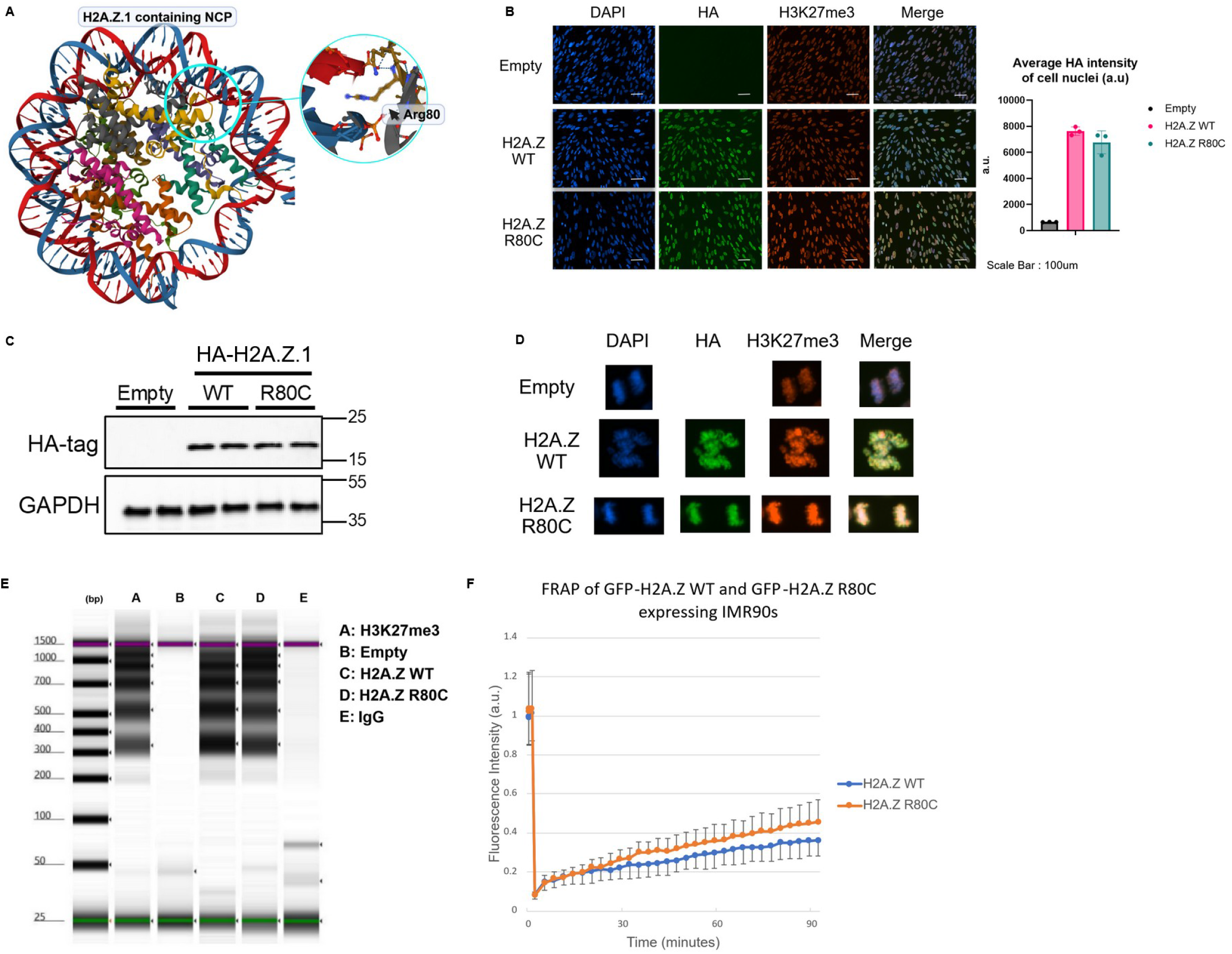
Biochemical characterization of wild type and mutants of histone variant H2A.Z in cells. (A) Location of H2A.Z R80 within the nucleosome core particle (NCP) (B) Representative images of cells co-stained with DAPI, anti-H3K27me3, and anti-HA antibody including quantification of HA signal intensity in cell nuclei (n=4) (C) Western blotting of cells ectopically expressing HA-H2A.Z R80 in proliferating IMR90s using an anti-HA antibody. (D) Representative images of dividing cells in anaphase showing the co-localization of DAPI, HA, and H3K27me3 signal from immunofluorescence. (E) High sensitivity DNA tapestation shows distinct nucleosomal ladder of ectopically expressed HA-H2A.Z histones in IMR90 cells (F) Fluorescence Recovery after Photobleaching (FRAP) was carried out in order to assess kinetics of histone mutants as measured by fluorescence recovery over 90 mins after photobleaching.

Several lines of evidence indicated that both H2A.Z WT and R80C are incorporated into nucleosomal chromatin. First, the fluorescent signal of the HA-tag co-localized with DAPI and H3K27me3 in both interphase and mitotic chromatin (Figure 1B, 1D). Co-localization of the WT and mutant histones on mitotic chromosomes strongly suggests incorporation of both variants into chromatin. A DNA digestion assay targeted by anti-HA antibodies using Tn5 transposase in IMR90 cells expressing either HA-tagged H2A.Z WT or R80C histones revealed clear nucleosomal ladder patterns, indicating that the ectopically expressed histones are both incorporated into chromatin (Figure 1E).

To assess the protein mobility of H2A.Z WT and R80C in the nucleus, we carried out fluorescence recovery after photobleaching (FRAP) to measure the kinetics of nuclear diffusion of GFP-tagged H2A.Z WT and R80C. The H2A.Z R80C mutant showed a more rapid fluorescence recovery rate compared to H2A.Z WT, indicating that H2A.Z R80C is more mobile within the nucleus, consistent with decreased nucleosome stability^40^ (Figure 1F). In sum, H2A.Z WT and R80C are comparably incorporated into nucleosomes and chromatin at steady state, but R80C is more mobile consistent with decreased nucleosome stability.

### H2A.Z R80C mutant suppresses induction of a subset of the SASP

Having characterized H2A.Z WT and R80C mutant in proliferating IMR-90 cells, we next investigated their impact on senescence-associated phenotypes. Upon irradiation, H2A.Z WT and H2A.Z R80C exhibited a comparable large and flattened morphology characteristic of senescence within 14 days (Figure 2A). Consistent with the induction of senescence, for both H2A.Z WT and R80C cells we observed a clear downregulation of genes associated with cellular proliferation and cell cycle progression following irradiation (Figure 2B). A principal component (PC) analysis of mRNA-seq at six time points, 0, 5, 8, 10, 12, and 14 days post-irradiation, revealed that roughly a quarter of transcriptomic variance (25.9%) was explained by PC1 with a clear separation of samples by time post-irradiation (Supplementary Figure 1). H2A.Z R80C tended to separate from empty and H2A.Z WT along PC2.

**Figure 2.**
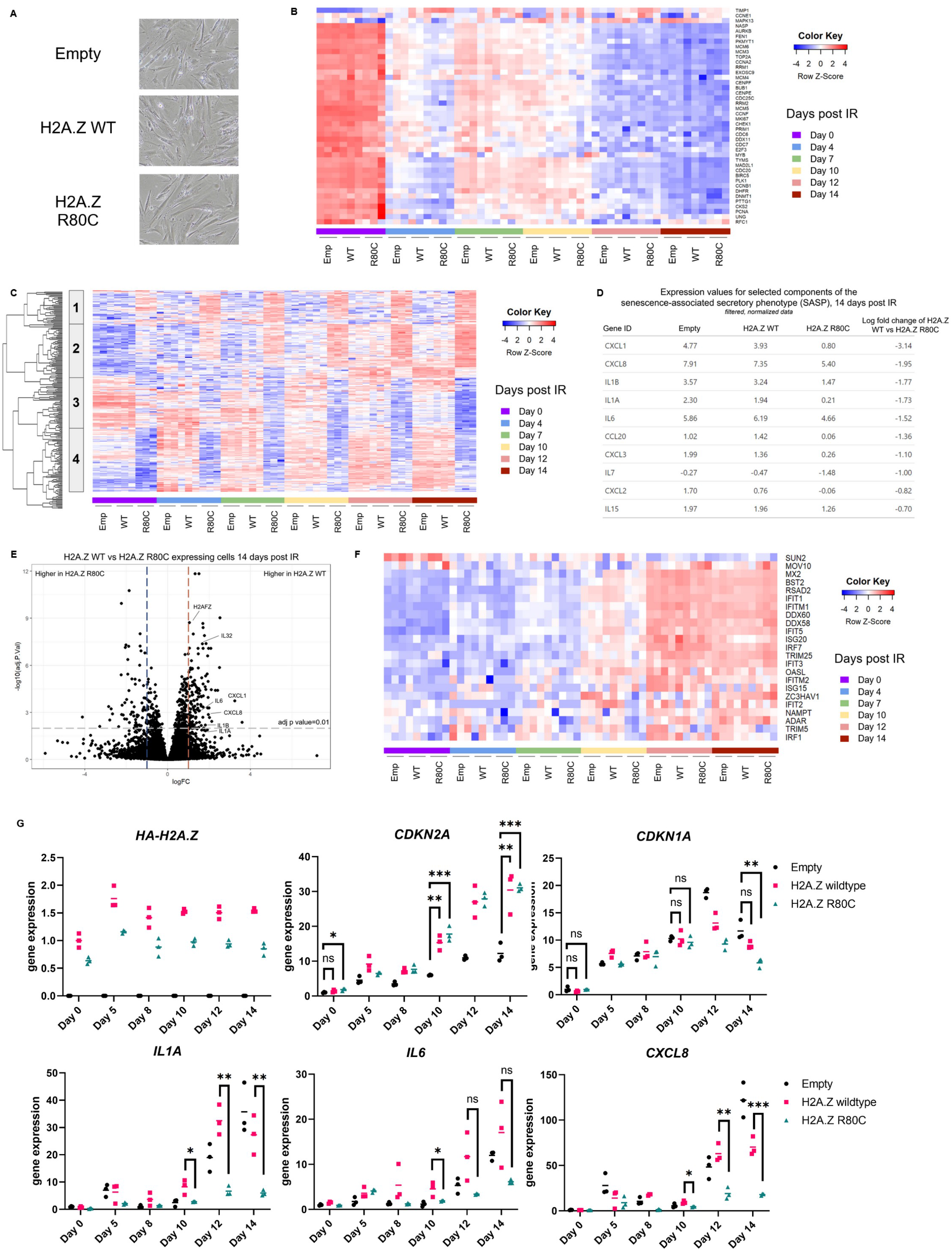
R80C histone variant H2A.Z specifically suppresses the SASP of senescent cells. (A) Images of senescent cells expressing H2A.Z WT and R80C mutant histone proteins (B) Heatmap showing the expression dynamics of genes associated with proliferation and cell cycle progression over a 14-day time course following irradiation (C) Heatmap of overlapping differentially expressed genes between H2A.Z WT vs H2A.Z R80C and Empty vs H2A.Z R80C expressing cells 14 days post IR. Gene signature clusters 1-4 are marked. (D) Expression and fold change of selected components of the SASP for senescent cells 14 days post irradiation (E) Volcano plot of H2A.Z WT vs H2A.Z R80C expressing cells at day 14 post irradiation. (F) A panel of interferon stimulated genes show broad upregulation during senescence but no significant changes in expression between H2A.Z WT and R80C mutant expressing cells (G) RT-qPCR of key SASP markers *IL1A, IL6*, and *IL8* show downregulation in H2A.Z R80C compared to H2A.Z WT expressing senescent cells without affecting expression of *CDKN1A* (p21) and *CDKN2A* (p16). Statistical significance was assessed using unpaired t-tests; \**P* < 0.05, \*\**P* < 0.01, \*\*\**P* < 0.001.

An unbiased clustering analysis of differentially expressed genes at day 14 post-irradiation identified four distinct gene clusters when comparing empty and H2A.Z WT controls and R80C-expressing cells (Figure 2C). Among these clusters, cluster 1 represented genes with expression levels that were unaffected during senescence in both empty and H2A.Z WT cells, but were upregulated in H2A.Z R80C cells. Cluster 2 included genes that were upregulated during senescence in empty, H2A.Z WT and H2A.Z R80C cells, albeit more so and less so in R80C in some cases. Cluster 3 contained genes downregulated during senescence in empty and H2A.Z WT cells, with further decrease in H2A.Z R80C. Finally, cluster 4 represented genes markedly upregulated during senescence in empty and H2A.Z WT cells, but less so in H2A.Z R80C cells.

Functional enrichment analysis of each gene cluster revealed distinct biological associations: cluster 1 was enriched with genes related to glycerolipid and cholesterol metabolism; cluster 2 with genes involved in cytokine and inflammatory responses; cluster 3 with genes related to extracellular matrix organization; and cluster 4 with genes encoding cell surface proteins (Supplementary Figure 2). Further analysis of these differentially expressed genes using the STRING database supports the results of this functional enrichment analysis in identifying both a group of pro-inflammatory genes and a cluster of collagen and matrix metalloproteinase genes that are downregulated in H2A.Z R80C expressing cells (Supplementary Figure 3).

We validated these functional enrichment analyses with a gene set enrichment analysis (GSEA) on the same set of differentially expressed genes and observed that expression of genes in pathways related to inflammation was repressed in H2A.Z R80C samples at day 14 post irradiation compared to H2A.Z WT (Supplementary Figure 4). Repression of genes related to inflammation was also apparent in parts of cluster 2 (IL1A, IL1B, IL6, CXCL3, CXCL5) and cluster 4 (CXCL1, CXCL8) (Supplementary Figure 2). These functional enrichment results indicate that there are two dominant repressive gene expression changes associated with expression of the H2A.Z R80C mutant during senescence, with one centered on the repression of a pro-inflammatory gene signature and the other on the repression of genes associated with the extracellular matrix.

Given that the SASP is enriched in both inflammatory cytokines/chemokines and extracellular matrix components, we examined whether the gene signatures from our unbiased analysis reflect the known SASP signature^13^. We examined the expression of several key SASP genes and found that their expression in senescence was lower in H2A.Z R80C expressing cells compared to H2A.Z WT (Figure 2D). Expression of these key SASP genes was significantly downregulated in senescent H2A.Z R80C expressing cells (Figure 2E). Importantly, the expression of H2A.Z R80C did not affect the expression of canonical interferon-stimulated genes (ISGs) that were upregulated during senescence (Figure 2F), confirming that H2A.Z R80C specifically affects a subset of the SASP. We verified suppression of key SASP markers *IL1A, IL6*, and *IL8/CXCL8* in H2A.Z R80C expressing cells by qPCR, while the expression of cell cycle genes *CDKN2A* and *CDKN1A* was not significantly impaired (Figure 2G). In summary, these results indicate that expression of H2A.Z R80C specifically suppresses genes associated with the SASP, including chemokines and cytokines involved in the proinflammatory phenotype of senescence.

### SASP suppression by H2A.Z R80C is specific to senescence and independent of loss of H2A.Z function

Next, we set out to investigate the mechanism by which H2A.Z R80C suppresses the SASP. First, we asked whether suppression of SASP is specific to the R80C substitution or can also be achieved by other amino-acid substitutions. To do this, we tested various H2A.Z R80 substitutions for their effect on senescence, including bulky residues (R80W), and those with altered (R80D, R80E) or similar (R80K) charge properties to the original arginine residue. Expression of all mutants was confirmed by RT-qPCR (Figure 3A). We observed similar reductions in SASP gene expression during senescence across all mutations (Figure 3B), indicating that SASP suppression is agnostic to the specific identity of the residue in the substitution. This is consistent with structural studies showing that R80 makes specific contact with DNA, and suggests that this interaction is disrupted by any mutation^41^.

**Figure 3.**
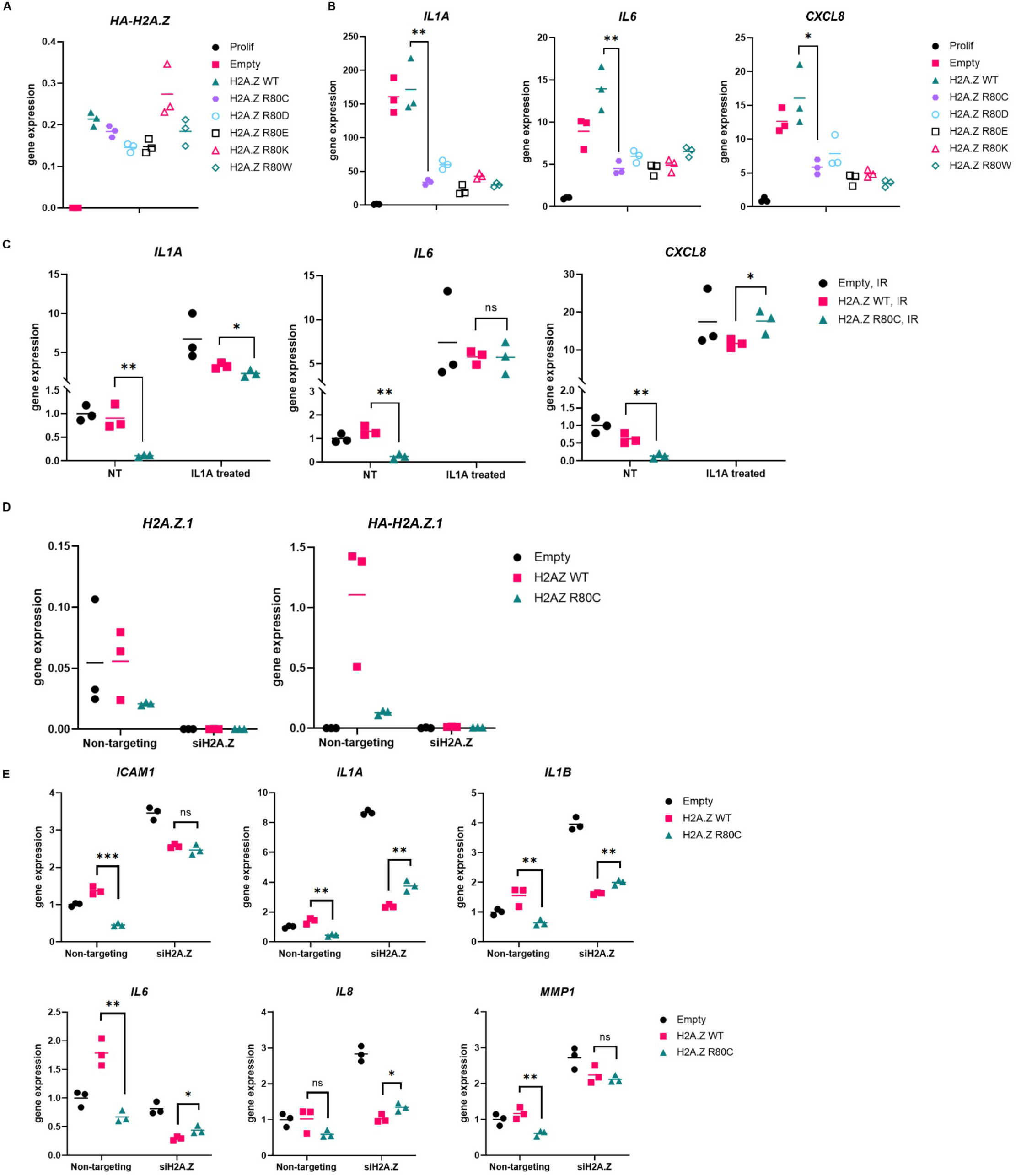
Mechanistic analysis of SASP suppression by R80C histone variant H2A.Z. (A) Changing R80 mutant does not notably change reduction of SASP in R80 mutant expressing senescent cells. (B) SASP gene activation at varying concentrations of IL1A compared to untreated cells (NT) in media does not differ significantly between H2A.Z WT and H2A.Z R80C expressing proliferating IMR90 cells. (C) SASP gene activation is not significantly different between H2A.Z WT and H2A.Z R80C expressing senescent IMR90 cells with exposure to 100pg/ml of IL1A for 6 hours in media in comparison to untreated cells (NT). (D) Validation of total H2A.Z and ectopically expressed HA-H2A.Z using RT-qPCR showed a highly efficient knockdown of H2A.Z histones. (E) Knockdown of H2A.Z abrogates reduction of SASP in H2A.Z R80C expressing compared to H2A.Z WT expressing senescent IMR90 cells. Statistical significance was assessed using unpaired t-tests for all panels; \**P* < 0.05, \*\**P* < 0.01, \*\*\**P* < 0.001.

Second, we asked whether H2A.Z R80C blocks expression of the canonical SASP genes only when they were induced as part of the senescence program. To investigate this, we added recombinant IL1A, a pro-inflammatory cytokine, into the media of senescent IMR90 cells and observed expression of SASP components *IL1A, IL6* and *IL8* by RT-qPCR. Under these conditions, we observed the reduction in *IL1A, IL6* and *IL8* expression by H2A.Z R80C was largely absent (Figure 3C). This indicates that H2A.Z R80C does not affect the capacity of cells to express *IL6* and *IL8* in response to other non-senescence triggers. Its suppressive effect on these SASP components is specific to the senescence program.

Finally, we wanted to know whether the suppression of the SASP by H2A.Z R80C is due to a loss of H2A.Z function. To test this, we asked whether suppression of the SASP was also achieved by H2A.Z knockdown. We performed siRNA knockdown of H2A.Z in senescent cells using a pool of three different siRNAs targeting H2A.Z.1. H2A.Z.2 (H2AFV) was not included in this analysis due to its substantially lower expression levels relative to H2A.Z.1 in our system, making it less likely to contribute significantly to the observed effects (Supplementary Figure 5). Using RT-qPCR, we verified knockdown efficiency of

98% based on transcript level of both total H2A.Z as well as ectopically expressed HA-tagged H2A.Z (Figure 3D). As expected, knockdown of ectopic H2A.Z R80C abrogated the reduction of the SASP by H2A.Z R80C, confirming that H2A.Z R80C in senescent cells represses the SASP (Figure 3E). However, knockdown of H2A.Z did not suppress the SASP gene expression in both Empty and H2A.Z WT cells. Indeed, knockdown of H2A.Z increased expression of SASP genes in several cases, suggesting a transcription repressive role in some cases. Regardless, these results indicate that the suppression of the SASP during senescence by H2A.Z R80C is not due to a loss of H2A.Z function.

### H2A.Z R80C shows altered epigenetic signatures in image-based deep learning classification

Given the role of histone modifications in control of gene expression, we hypothesized that the altered transcriptomic program of H2A.Z R80C-expressing cells is linked to changes in epigenetic signatures. We employed deep learning techniques to examine differences in various epigenetic signatures in proliferating and senescent IMR-90 cells using immunofluorescent images of key regulatory histone modifications: H3K27ac, H3K27me3, H3K4me1, and H3K56ac^42,43^. Comparisons among different neural networks revealed that the ResNet50 pre-trained convolutional neural networks performed best to differentiate between proliferating cells based on immunofluorescence staining of H3K27ac, H3K27me3, H3K4me1, H3K56ac and DAPI staining (Figure 4A). The trained ResNet50 model robustly distinguished various nuclear epigenetic marks, while highlighting a visually similar signature between the enhancer marks H3K27ac and H3K4me1 compared to other modifications, demonstrating the power of this machine learning approach to detect biological differences in epigenetic signatures between individual cells (Figure 4B). We observed that, based solely on DAPI staining, a separate trained ResNet model was able to robustly distinguish between proliferating and senescent cells (Figure 4C). While DAPI alone could not clearly distinguish R80C cells from others, H3K27ac, H3K27me3, and H3K4me1 clearly separated it from both Empty and H2A.Z WT cells (Figure 4D). These results suggest that specific chromatin marks such as H3K27ac and H3K27me3 reflect distinct epigenetic states between H2A.Z WT and R80C-expressing cells.

**Figure 4.**
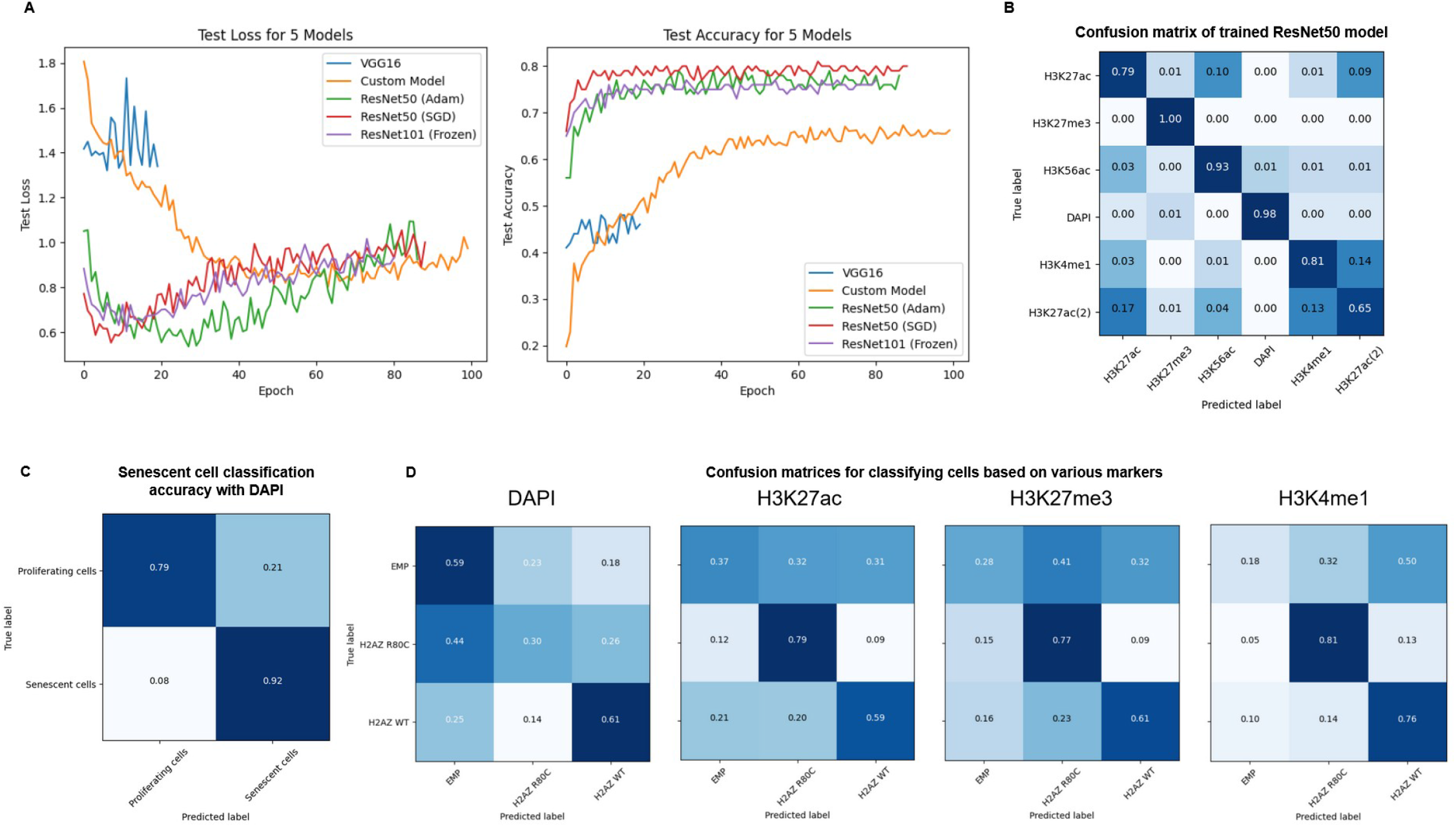
Image classification using deep learning models to investigate consequences of wild type and mutant histone variant H2A.Z. (A) Binary Cross Entropy Loss and Accuracy on test data during training of different model architectures after training for 80 to 100 epochs (B) Cells stained with different epigenetic markers can be robustly classified using a trained deep learning model (C) A trained ResNet101 was able to robustly classify proliferating and senescent cells based solely on images of DAPI staining of cell nuclei (D) Cells ectopically expressing H2A.Z R80C show distinct epigenetic patterns that can distinguished by deep learning models.

### Epigenetic signatures at SASP genes are altered by H2A.Z R80C expression

Based on this finding, we wondered whether the repression of the SASP genes by H2A.Z R80C might be associated with altered histone modifications at those genes. To test this, we performed CUT&Tag for histone modifications H3K27ac, H3K27me3 and H3K4me1 as well as for HA-tagged H2A.Z WT and R80C in proliferating and senescent cells. Clustering of samples based on genome-wide distribution of HA-tagged H2A.Z wild type and R80C mutant showed greater correlation with distribution of H3K27ac than with H3K27me3, consistent with preferential enrichment of H2A.Z at active gene loci (Figure 5A). The enrichment level of H2A.Z R80C did not significantly differ from that of H2A.Z WT across proliferating and senescent cells in H2A.Z WT and R80C expressing cells (Supplementary Figure 6). By contrast, H3K27ac, an activating histone mark, was significantly enriched at differentially expressed SASP gene loci during senescence, but this was not the case in H2A.Z R80C expressing cells (Figure 5B). These findings suggest that H2A.Z R80C may suppress SASP gene expression by attenuating recruitment of activating H3K27ac. Together, these results demonstrate that H3K27ac enrichment at SASP gene loci depends on the stability of H2A.Z. These findings collectively suggest that H2A.Z R80C modulates SASP gene expression by interfering with H3K27ac-associated chromatin activation.

**Figure 5.**
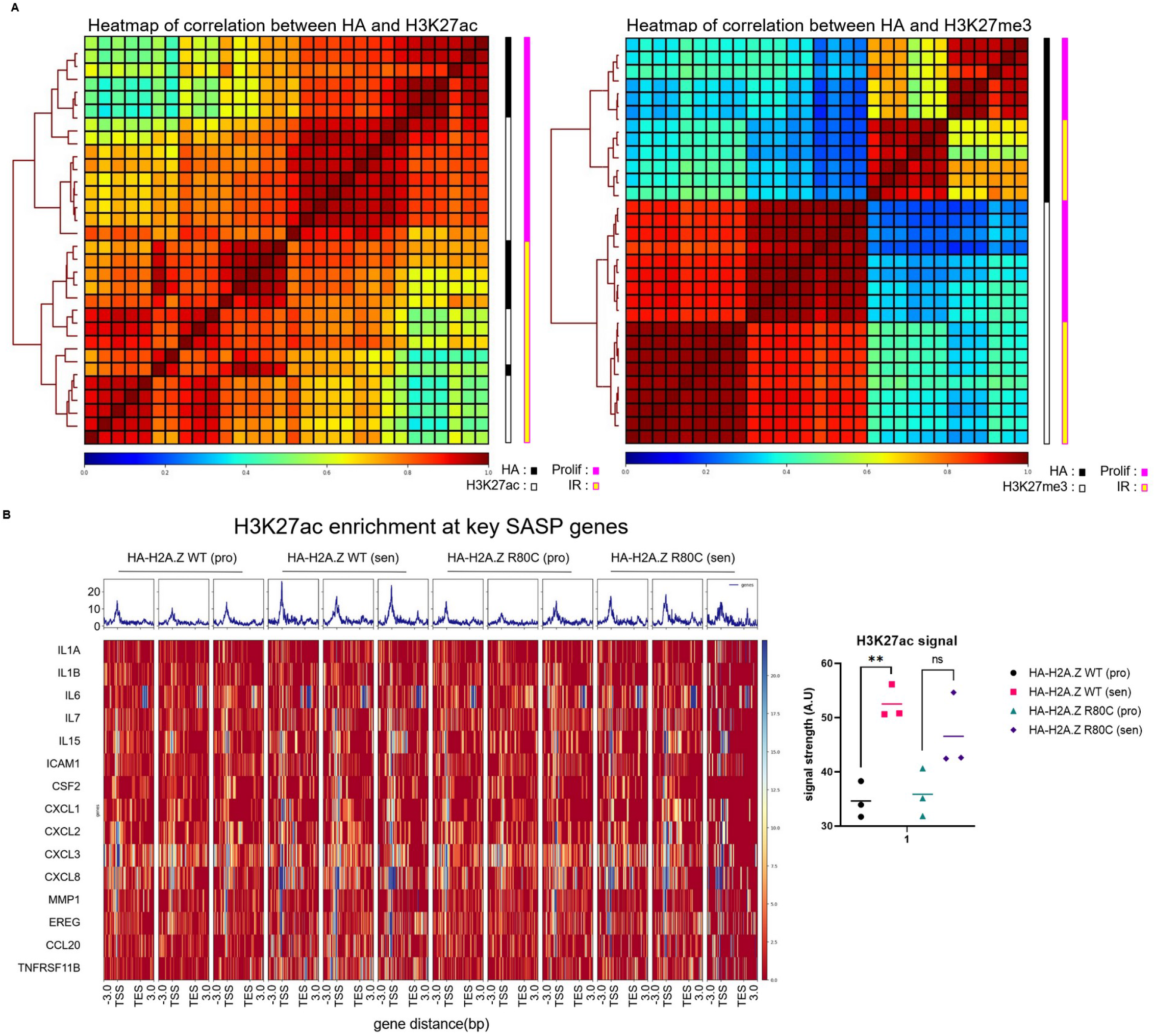
Epigenome distribution of wild type and mutant histone variant H2A.Z. (A) Heatmap of signal correlation between HA tag and H3K27ac and HA tag and H3K27me3 signal between proliferating and senescent conditions of IMR90 cells expressing both H2A.Z wild type and mutant R80C histones. (B) Enrichment at key SASP genes of HA tag and H3K27ac signal as assessed using CUT&TAG of senescent IMR90 cells expressing wild type and mutant H2A.Z histones at 11 days post irradiation. Statistical significance was assessed using unpaired t-tests; \**P* < 0.05, \*\**P* < 0.01, \*\*\**P* < 0.001.

## DISCUSSION

Our results reveal a novel role of the stability of H2A.Z-containing nucleosomes in regulating expression of the SASP during cellular senescence. Expression of the H2A.Z R80C mutant decreased the stability of the nucleosome core particle and specifically downregulated the SASP during senescence, without affecting upregulation of other genes in senescence, such as ISGs, *CDKN1A* and *CDKN2A*. The effect of H2A.Z R80C mutation was recapitulated by other substitutions of R80, was specific to senescence over other inducers of cytokine/chemokine expression, and was not due to loss of H2A.Z function. Rather, the effect of H2A.Z R80C appears to be due to impaired marking of SASP genes by H3K27ac.

H2A.Z R80C was first identified as a histone mutant that is enriched in cancer. While many histone mutants are seemingly random in nature (likely ‘passenger’ mutations), a small subset of them are recurrent and over-represented within cancer genomes, and are therefore considered potential ‘drivers’ of disease^40,44^ Such ‘oncohistone’ mutations are predicted to play a crucial role in affecting nucleosome structure and diverse cellular processes such as chromatin accessibility, DNA damage repair, replication, and histone post-translational modifications - ultimately predisposing cell fate towards transformation. Given the tumor suppressive role of the SASP in senescent cells through its ability to recruit the innate immune system, it is tempting to speculate that the oncogenic effect of H2A.Z R80C is, at least in part, due to suppression of the SASP^45^.

Our biochemical analysis of cells ectopically expressing either wild type or mutant H2A.Z proteins suggests that these ectopically expressed histones are incorporated into chromatin and nucleosomes in a manner that is similar to endogenous histones based on imaging and CUT&Tag data. We further observed through bulk RNA-seq that a subset of key SASP genes was repressed when cells ectopically express H2A.Z R80C. This included genes associated with the pro-inflammatory SASP, including *IL1A, IL1B, IL6, CXCL1, CXCL3, CXCL5*, and *CXCL8*^46^. However expression of ISGs which represent a separate arm of the senescence program that is distinct from the SASP as well as cell cycle genes *CDKN1A* and *CDKN2A* was not affected^47,48^. Notably numerous genes associated with the extracellular matrix and matrix metalloproteinases were downregulated during senescence and were further repressed in H2A.Z R80C-expressing cells, including *COL3A1, COL6A1, COL6A2*, and *PCOLCE*. This suggests that H2A.Z R80C selectively modulates senescence-associated genes that influence extracellular environment, rather than globally altering the senescence transcriptional program.

We have observed a similar phenotype in cells expressing different substitutions at the R80 site, specifically cysteine (C), aspartic acid (D), glutamic acid (E), lysine (K) and tryptophan (W). These results indicate that the arginine residue at H2A.Z R80 is essential for establishing the SASP phenotype. The residue R80 in histone H2A.Z plays an important role in influencing the structure and stability of the nucleosome due to its close interaction with the adjacent DNA strand that wraps around the nucleosome. Crystal structure modeling of the nucleosome including the H2A.Z histone variant indicates that the positively charged H2A.Z R80 arginine residue closely fits the minor groove of the adjacent negatively charged DNA strand^25^. This suggests that the R80 residue is important in modulating histone-DNA interactions within the assembled nucleosome, and that mutating this residue would potentially impact the kinetics of the formation/disassembly of H2A.Z-containing nucleosomes^41,49^. Results reported here indicate that substitution of R80 with any amino-acid is sufficient to disrupt its function. Interestingly, substitution of cysteine for arginine can occur through a simple C to T transition at the first base of the DNA codon, potentially through deamination of methylated c^me^GT (arginine) to TGT (cysteine) in H2A.Z.1. Together, these results and observations suggest that the frequency of R80C over other substitutions is in part a function of the genetic code, not a specific advantage for cancer of cysteine over substitutions.

We further tested if the suppression of the SASP by H2A.Z R80C was specific to the context of cellular senescence or if this mutant would also suppress the same cytokine/chemokines induced by a different trigger. We observed that, upon stimulation of pro-inflammatory genes with IL1A supplementation in media, the activation of genes was comparable between H2A.Z WT and R80C expressing cells, indicating the effect of R80C is specific to the context of senescence^50^.

Notably, knockdown of H2A.Z did not recapitulate the suppression of the SASP observed with R80C expression, suggesting that the effect is independent of H2A.Z abundance. This rules out a simple loss-of-function model where the R80C mutation merely leads to functional inactivation of H2A.Z. Instead, our data supports a model where the presence and incorporation of the destabilized H2A.Z R80C protein is required to actively suppress the SASP. Potentially, the mutant protein functions in a dominant-negative or neomorphic fashion.

The induction of the SASP is accompanied by redistribution of histone modifications to establish a pro-inflammatory gene expression program. We observed distinct visual epigenetic signatures in senescent cells expressing the R80C mutant using an unbiased, deep-learning approach analyzing immunofluorescent images. Models trained on specific histone modifications—particularly the active enhancer and promoter mark H3K27ac—could successfully discriminate between the two cell states. This initial finding pointed to a defect in the establishment of active chromatin in senescent cells. We confirmed this directly at target gene loci using CUT&Tag. During senescence, H2A.Z WT cells exhibited a significant enrichment of H3K27ac at SASP gene loci, consistent with their robust transcriptional activation. In stark contrast, this gain of H3K27ac was notably reduced in H2A.Z R80C-expressing cells.

In conclusion, we have observed a previously unremarked relationship between nucleosome stability and senescence phenotype. Decreasing the stability of H2A.Z containing nucleosomes results in a significant and specific reduction in the expression of pro-inflammatory SASP genes. This finding advances our understanding of cellular senescence, and how it can be modulated. More remains to be explored, especially in terms of the mechanism underlying this phenomenon and its relevance to aging biology. Modulating nucleosome stability may offer a strategy to intervene in senescence and pave the way for new therapeutics in the future that ameliorate the detrimental effects of cellular senescence.

## METHODS

### Cell culture

IMR90 fibroblasts and HEK293T cells were cultured in growth media DMEM (1x) (10% Fetal Bovine Serum, 2mM L-glutamine, 1% pen-strep) (Gibco cat # 10313021) and incubated at 37°C, 5% CO2, 3.5% 02 for the former and 37°C, 5% CO2 for the latter. IMR90s and HEK293Ts were split every 2 to 3 days or when confluent by incubating with 0.25% Trypsin-EDTA (Gibco cat # 25200056) for 5 mins within the cell incubator to detach adherent cells. A minimum of 3x volume of growth media is added to detached cells before cells are spun down at 300g for 3 minutes. Liquid above the cell pellet is then aspirated before the cell pellet is resuspended with growth media with gentle pipetting. Resuspended cells are replated on appropriate surface-treated polystyrene.

Frozen cell stocks are made from freezing down growing cells by pelleting cells as described above. Cell pellets are then resuspended in Fetal Bovine Serum with 10% DMSO. Resuspended cells are placed in cryovials in a styrofoam box with the lid closed in -80°C freezer for at least 6 hours. Cells are transferred to a liquid nitrogen tank afterwards for long term storage.

### Senescence induction

IR induced senescence was carried out by irradiating cells at a dose of 20Gy before an immediate refresh of the appropriate cell media. Irradiation was carried out within an X Ray lrradiator machine (Rad Source Technologies RS 2000 Small Animal lrradiator) that is regularly calibrated. Cells were split 1:3 three days post irradiation, and media of cells are refreshed every 2 to 3 days after becoming senescent and have stopped proliferating.

### Plasmid preparation

DNA sequences for the sequences encoding H2A.Z WT, H2A.Z R80C, H2A.Z R80D, H2A.Z R80E, H2A.K R80K, and H2A.Z R80W were ordered from Integrated DNA Technologies (IDT). pLenti-PGK plasmids used for subsequent lentivirus preparation were obtained from a collaborator and the plasmid sequence checked using Sanger sequencing.

Cloning of DNA sequences into plasmid was carried out using the restriction endonucleases BamHI (R3136S) and Sall (R3138S) for digestion of both insert and plasmid sequences while T4 ligase (M0202L) was used for the subsequent ligation reaction. All enzymes used were obtained from New England Biolabs and utilized based on recommended incubation time and temperature.

Cloned plasmids were transformed into stable competent C3040 E. coli using heat shock for one minute at 42°C and then immediately chilling the sample on ice before streaking on sterile Agar plates containing Carbenicillin for selection of transformed E. coli. After incubation of plate for 24 hours at 37°C single colonies are picked using a sterile pipette tip and grown in Luria-Bertani **(LB)** broth for 12 hours. **LB** broth was prepared with LB medium capsules (3002011) from MP Biomedicals and ultrapure water from a Barnstead™ GenPure™ xCAD Plus Ultrapure Water Purification System (Thermo Fisher Scientific, cat # 50136151). Plasmid DNA was subsequently isolated using a ZymoPURE II Plasmid Midiprep Kit (Zymo Research, cat # D4201) and sequence verified using Sanger sequencing.

### Lentivirus generation and use

HEK293T cells were grown to 50-60% confluence and transfected using Lipofectamine 2000 (Thermo Fisher Scientific, cat # 11668030) with 2.4ug/ml of lentiviral expression vector, 1ug/ml psPAX2 packaging plasmid and 0.6ug/ml VSVG plasmid over 6 hrs. Viral laden media was collected after 24 hrs post transfection and aliquoted and stored at -80 C for long term storage.

Cells were infected by lentivirus by thawing frozen viral laden media and adding it to fresh growth media of cells after titrating viral MOI. After 24 hours have passed media is refreshed with fresh growth media containing 1ug/ml of puromycin for selection. After 72 hours have passed, cells are split and replated with fresh growth media.

### Antibodies

For western blots and CUT&TAG against GFP we used Abeam anti-GFP antibody (ab6556), while anti-H3K27ac (ab177178) from Abeam and anti-H3K27me3 (9733S) from Cell Signaling Technology was used for CUT&TAG. We further used antibodies against alpha tubulin from Santa Cruz (sc-8035), actin from Sigma-Aldrich (A1978) and H2A.Z from Cell Signaling Technology (2718S) in western blots.

For CUT&TAG and immunofluorescence staining against specific epigenetic markers for deep learning based epigenetic analysis, we used the antibodies anti-H3K56ac from Active Motif (39281), anti-H3K4me1 from Abeam (ab8895), anti-H3K27ac from Abeam (ab4729) and from Active Motif (39133), anti-H3K27me3 from Cell Signaling (9733S), anti-H3K4me2 from Sigma-Aldrich (07-030). We used as an IgG control, antibodies from Vector Laboratories (I-1000, I-2000).

We used the antibodies against FLAG tag from Cell Signaling Technology (14793S) and against HA tag from Santa Cruz (sc-7392) for western blotting, CUT&TAG, and immunofluorescence staining.

### Western Blot

Proteins are extracted from cells using sample buffer (0.5M TrisHCI ph6.8, Glycerol, SDS, BPB) and ran in denaturing gels. Proteins are blotted on to lmmobilon-P PVDF Membrane (MiliporeSigma cat # IPVH00010) using BioRad transblot Turbo (BioRad cat # 1704150) before staining with Ponceau S (Sigma-Aldrich cat # P7170S-1L). Membrane is blocked at room temperature for 30 mins in TBST (Tris-buffered saline, 0.1% Tween 20) with 5% milk and left on a shaker in the dark overnight at 4C with antibody targeting protein of interest. After 3× 10 min washes in TBST, secondary antibodies are diluted to 1:10000 in 5% milk with TBST and incubated with membrane for 1hr at room temperature. Protein bands are imaged using SuperSignal™ West Femto Maximum Sensitivity Substrate (Thermo Fisher Scientific, cat # 34094).

An alternative Western Blot protocol using Bullet Blocking buffer (Nacalai USA cat # 13779-01) instead of 5% milk was also employed to achieve similar results.

### RNA-seq

RNA-seq was carried out on the Illumina platform and all analysis was performed in R. Alignment was carried out using pseudo-alignment with Kallisto against hg38 cDNA. R packages used include tidyverse, edgeR, and matrixStats for statistical calculations, as well as plotly, ggrepel, gt, ggplot, gplots, RColorBrewer, and enrichplot, for tables, heatmaps, and plots. GSEA and GO term analysis was carried out in R using the packages GSEABase, msigdbr, and gprofiler2

### lmmunofluorescence

Cells were washed with DPBS (Gibco, cat # 14190144) before fixing with 10% Neutral Buffered Formalin (Thermo Fisher Scientific, cat # 9400-1) for 10 mins at room temperature. Cells were incubated in blocking solution (4% bovine serum albumin, 1% goat serum in DPBS) before antibody staining in blocking solution according to the manufacturer’s recommended concentration of antibodies. Secondary antibodies used are Alexa Fluor Goat anti-Rabbit IgG 488 (A11008) and Goat anti-Mouse (A11029) as well as lnvitrogen Goat anti-Rabbit IgG 555 (A32732) and Goat anti-Mouse 555 (A32727). DAPI staining was carried out using DAPI solution (100ng/ml DAPI in DPBS).

### RNA quantification

After a DPBS wash step RNA is extracted directly from cells through addition of RNA lysis buffer on to plate and proceeding with steps outlined in Quick-RNA MiniPrep kit (Zymo Research, cat # 11-328). RNA is reverse transcribed using ReverAid RT (Thermo Fisher Scientific, cat # K1691) to obtain cDNA which is diluted 1:10 in water and quantified in 384 well format with SYBR™ Green PCR Master Mix (Thermo Fisher Scientific, cat # 4309155) with appropriate primers (full list in Supplementary Table 1).

### siRNA knockdown

siRNA knockdown was performed using a 20nM pool of three different siRNAs targeting H2A.Z using the Trifect RNAi hs.Ri.H2AFZ.13 by IDT. Cells were treated with 3ul RNAiMAX (Thermo Fisher Scientific, cat #13778150) per 6 well volume along with pooled siRNA for 24hr before media was refreshed with fresh growth media. Knockdown is measured at least 48 hrs after treatment of cells with siRNA.

### CUT&TAG

CUT&TAG was carried out according to the original protocol outlined online by Steven Henikoff (https://www.protocols.io/view/cut-amp-tag-home-bd26i8he), using native nuclei isolated from proliferating IMR90 cells. 100,000 nuclei were used for each sample. Single amendment was made to use Halt™ Protease Inhibitor Cocktail (100X) (cat # 78430) instead of Roche Complete Protease Inhibitor EDTA-Free tablets. CUT&TAG pAG-Tn5 protein was from Epicypher (SKU: 15-1017).

Analysis was carried out using bowtie2 for alignment and SEACR for peak calling. Peak annotation was carried out using annotatePeaks.pl from HOMER. Alignment and annotations were carried out using the latest version of hg38. Heatmaps were generated using deepTools.

### Epigenetic characterization with deep learning

All algorithms and model training were developed in Jupyter with Python 3.10. Pre-trained ResNet models were downloaded using Pytorch and weights were frozen before unfreezing the last layer during training.

All code used to carry out cell segmentation, training of ResNet models, and classification is publicly available on GitHub (https://github.com/zongmingchua/Epigenetic_image_ML)

## DATA AVAILABILITY

The sequencing datasets generated and analyzed during the current study are not yet publicly available due to ongoing processing in the Gene Expression Omnibus repository but will be available from the corresponding author on reasonable request or upon publication of this manuscript.

## AUTHORS CONTRIBUTIONS

P.O. A conceived the concept of the study, procured the funding, and was in charge of the overall direction. Z.M and H.T produced the results shown in Fig 1. Z.M produced the results shown in Figs 2-5. A.A assisted in producing plasmids encoding different substitution mutants at the H2A.Z R80 site. A.R and X.L assisted with advice in conducting bioinformatics and machine learning analysis. M.G.T, K.M, A.D, C.C.M, L.H, and N.D assisted with refining experimental protocols and general technical analysis. All authors including were involved in the preparation of the manuscript.

