## Supplementary Material for "Histone variant H2A.Z mutant suppresses the senescence-associated secretory phenotype"

Supplementary Table 1
List of Sybr green primers used for qPCR

| Gene Name | Forward and Reverse PCR primer sequence |
|---|---|
| *ICAM1* | GGTTGAACCCCACAGTCACCTATG<br>GCTGTAGATGGTCACTGTCTGCAG |
| *IL1A* | AGTGCTGCTGAAGGAGATGCCTGA<br>CCCCTGCCAAGCACACCCAGTA |
| *IL1B* | TGCACGCTCCGGGACTCACA<br>CATGGAGAACACCACTTGTTGCTCC |
| *IL6* | AAATTCGGTACATCCTCGACG<br>TTTCACCAGGCAAGTCTCC |
| *IL8* | GAGTGGACCACACTGCGCCA<br>TCCACAACCCTCTGCACCCAGT |
| *MMP1* | ATCGGCCCACAAACCCCAAA<br>TGGCAGTTGTGGCCAGAAAACA |
| *ACTB* | AGAGCTACGAGCTGCCTGAC<br>AGCACTGTGTTGGCGTACAG |
| *GAPDH* | GAAGGTGAAGGTCGGAGTC<br>TTGAGGTCAATGAAGGGG |
| *CDKN2A* | CTGCCCAACGCACCGAATAG<br>CCACCAGCGTGTCCAGGAAG |
| *CDKN1A* | CTGCAGGGGACAGCAGAGG<br>CCGGCGTTTGGAGTGGTAG |
| *HA-H2A.Z* | ATGGCCTACCCCTACGACGTG<br>AACCGCCTTTGTCTTGGCCTTTC |
| *H2AZ1* | GGTCCGATTAGCCTTTTCTCTG<br>AACCGCCTTTGTCTTGGCCTTTC |

Supplementary Figure 1

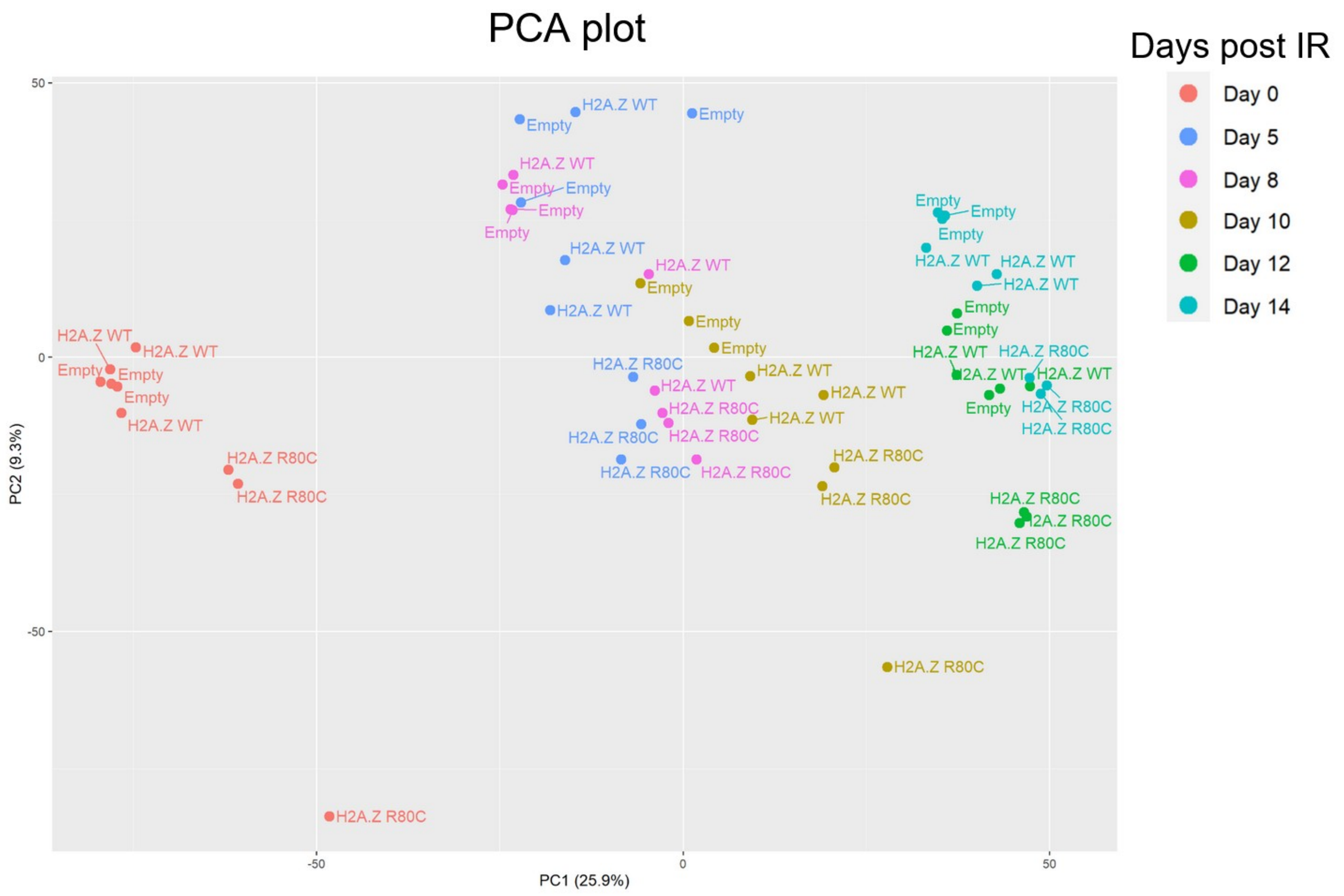


PCA plot showing PC1 and PC2 of bulk RNA-seq samples colored based on days elapsed post irradiation, including percentage of variances explained by the respective primary component

Supplementary Figure 2

### Heatmap of differentially expressed genes in cluster 1

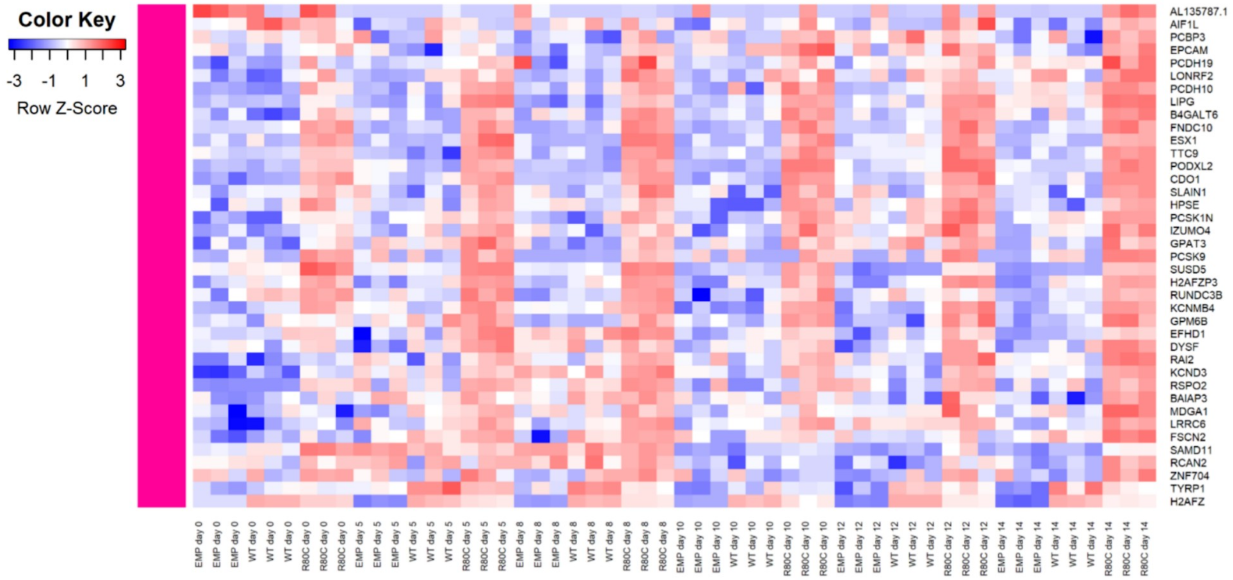


### Functional enrichment analysis of cluster 1

| id | source | term_id | term_name | term_size | | p_value |
|---|---|---|---|---|---|---|
| 1 | KEGG | KEGG:00561 | Glycerolipid metabolism | 61 | | 4.2e-02 |
| 2 | KEGG | KEGG:04979 | Cholesterol metabolism | 50 | | 4.2e-02 |
| 3 | WP | WP:WP4522 | Metabolic pathway of LDL, HDL and TG, including diseases | 17 | | 3.4e-03 |

*g:Profiler (biit.cs.ut.ee/gprofiler)*

### Heatmap of differentially expressed genes in cluster 2

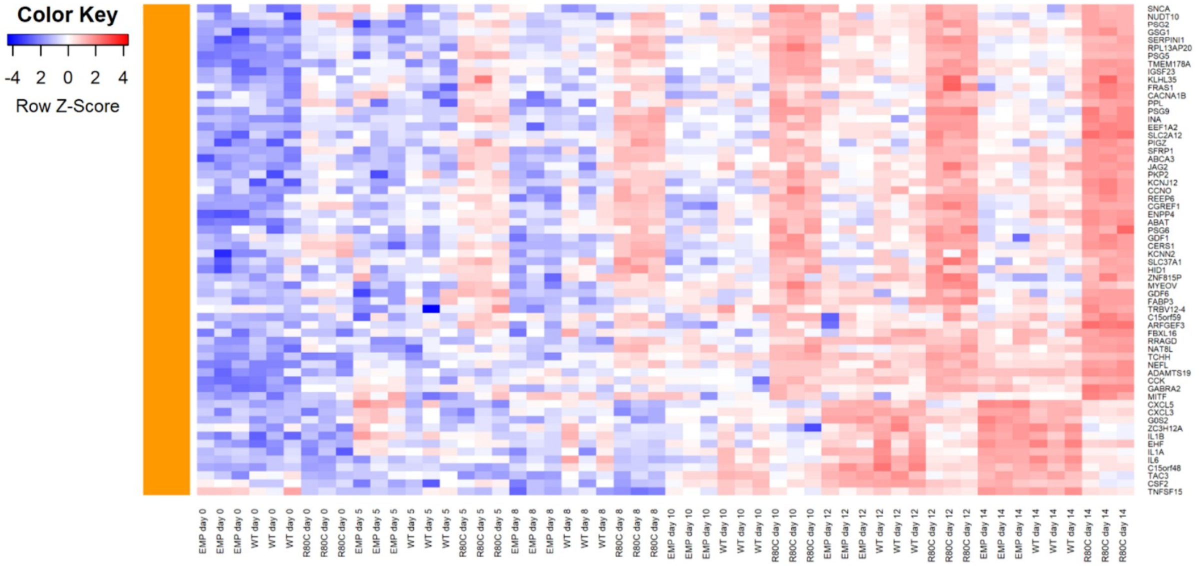


### Functional enrichment analysis of cluster 2

| id | source | term_id | term_name | term_size | | p_value |
|---|---|---|---|---|---|---|
| 1 | GO:MF | GO:0005125 | cytokine activity | 235 | | 5.1e-06 |
| 2 | GO:MF | GO:0030546 | signaling receptor activator activity | 502 | | 1.2e-05 |
| 3 | KEGG | KEGG:04060 | Cytokine-cytokine receptor interaction | 293 | | 5.9e-05 |
| 4 | WP | WP:WP530 | Cytokines and inflammatory response | 26 | | 3.7e-04 |
| 5 | WP | WP:WP5095 | Overview of proinflammatory and profibrotic mediators | 127 | | 4.2e-04 |

*g:Profiler (biit.cs.ut.ee/gprofiler)*

### Heatmap of differentially expressed genes in cluster 3

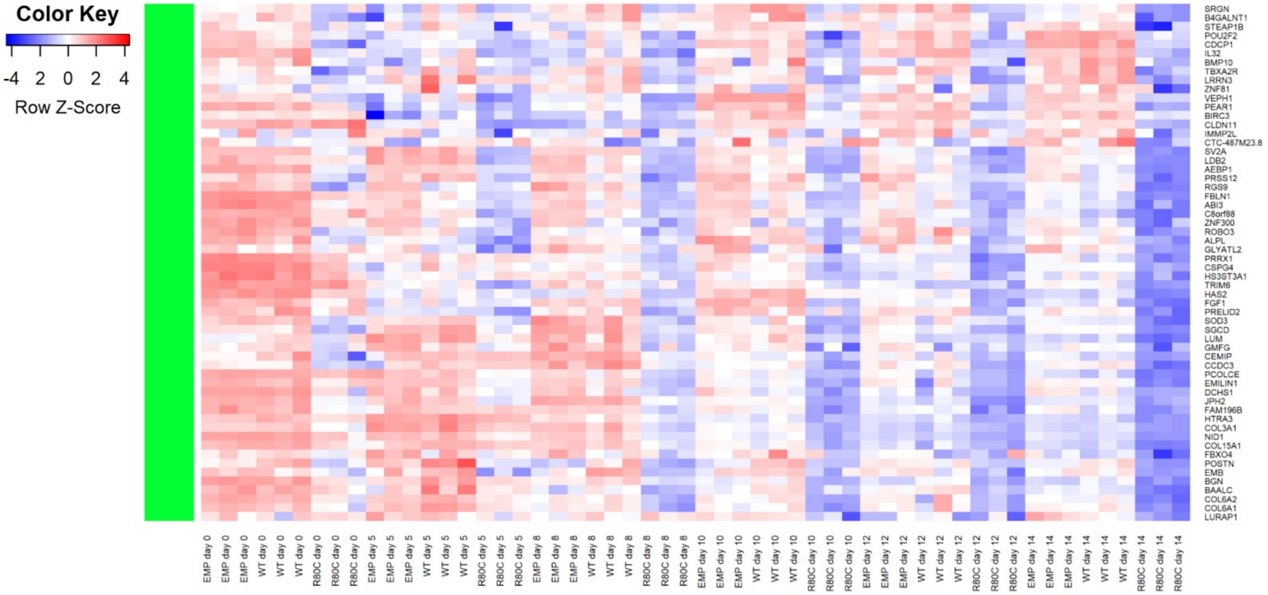


### Functional enrichment analysis of cluster 3

| id | source | term_id | term_name | term_size | | p_value |
|---|---|---|---|---|---|---|
| 1 | GO:MF | GO:0005201 | extracellular matrix structural constituent | 173 | | 5.8e-12 |
| 2 | GO:MF | GO:0005518 | collagen binding | 69 | | 3.1e-06 |
| 3 | GO:CC | GO:0031012 | extracellular matrix | 560 | | 3.6e-12 |
| 4 | REAC | REAC:R-HSA-1474244 | Extracellular matrix organization | 298 | | 1.5e-07 |
| 5 | HPA | HPA:0461432 | skin 1; extracellular matrix[≥Medium] | 48 | | 6.6e-12 |

*g:Profiler (biit.cs.ut.ee/gprofiler)*

### Heatmap of differentially expressed genes in cluster 4

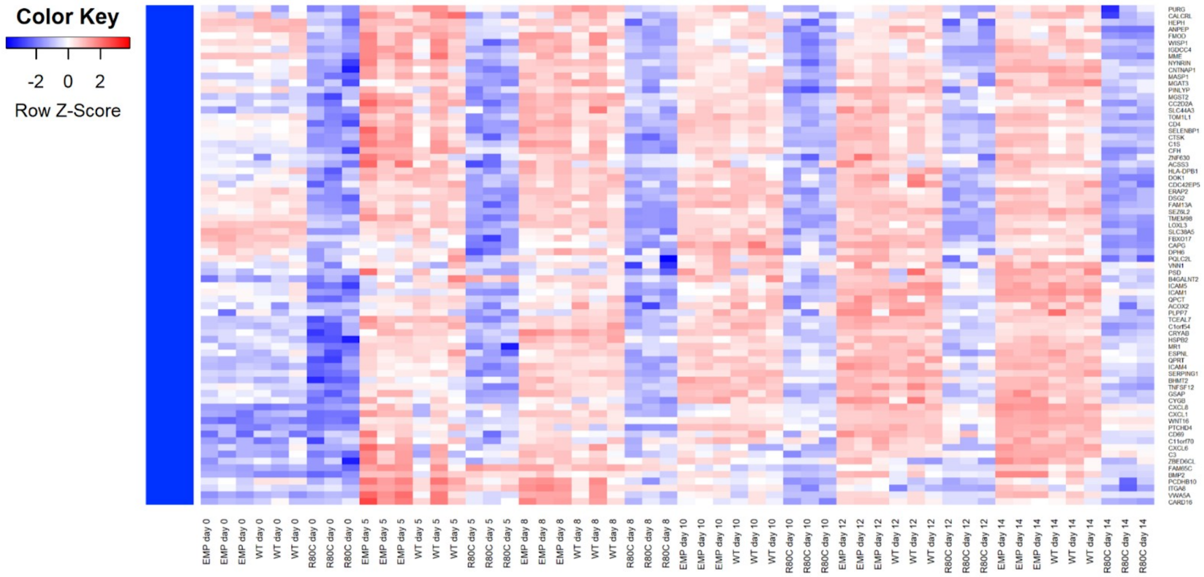


### Functional enrichment analysis of cluster 4

| id | source | term_id | term_name | term_size | | p_value |
|---|---|---|---|---|---|---|
| 1 | GO:CC | GO:0005576 | extracellular region | 4303 | | 2.6e-04 |
| 2 | GO:CC | GO:0009986 | cell surface | 908 | | 4.2e-04 |
| 3 | WP | WP:WP2806 | Complement system | 99 | | 3.6e-04 |

*g:Profiler (biit.cs.ut.ee/gprofiler)*

### Supplementary Figure 3

Local network cluster (STRING)

| Cluster | Count in Network | Strength | False Discovery Rate | Description |
|---|---|---|---|---|
| CL:18273 | 6 of 34 | 1.38 | 0.00047 | Chemokine receptor binding |
| CL:16433 | 6 of 57 | 1.16 | 0.0032 | Collagen biosynthesis and modifying enzymes |
| CL:16569 | 6 of 58 | 1.15 | 0.0032 | Mixed, incl. elastic fibre formation and MMP |

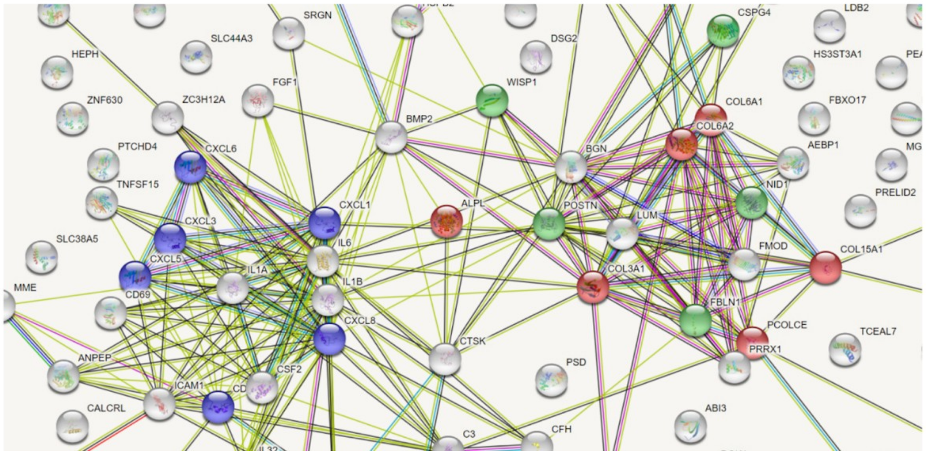


### Supplementary Figure 4

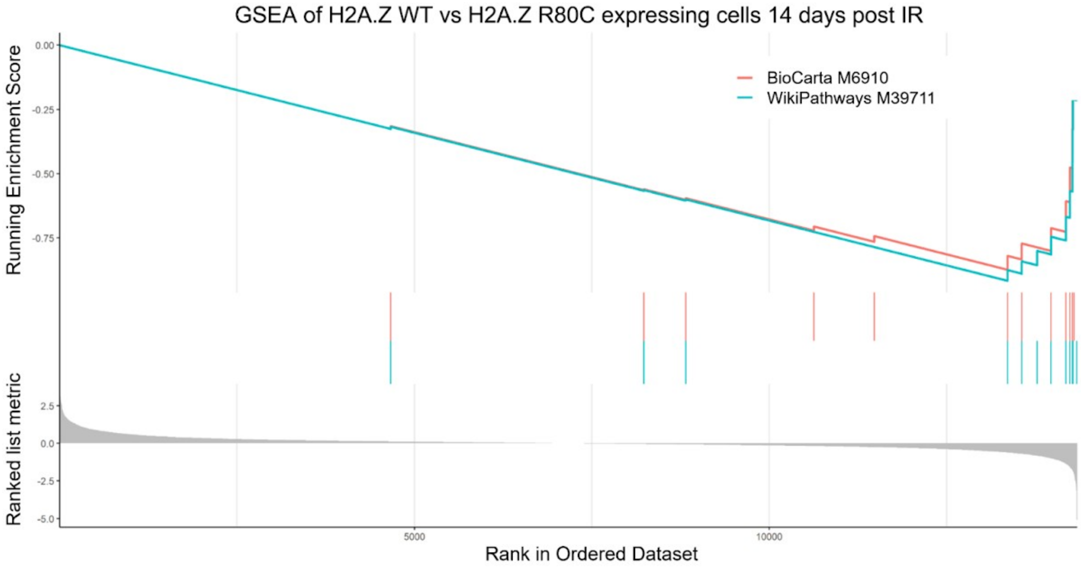


(A) STRING database analysis of genes enriched in H2A.Z WT expressing cells compared to H2A.Z R80C expressing cells at 14 days post irradiation (B) Gene Set Enrichment Analysis shows that genes associated with Cytokines and Inflammatory Response (BioCarta M6910 (BIOCARTA_INFLAM_PATHWAY) and WikiPathways M39711 (WP_CYTOKINES_AND_INFLAMMATORY_RESPONSE)) are strongly enriched in H2A.Z WT expressing cells compared to H2A.Z R80C expressing cells. BioCarta M6910 has an enrichment score of -0.87 and WikiPathways M39711 an enrichment score of -0.92.

Supplementary Figure 5

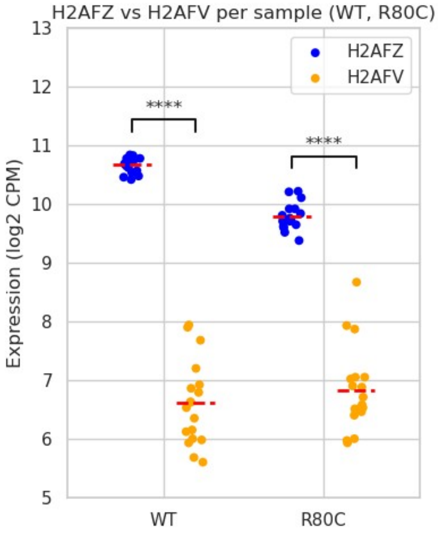


Expression (log2 CPM) of H2AFV and H2AFZ as observed in H2A.Z WT and H2A.Z R80C expressing cells from days 0 to 14 during senescence induction.

Supplementary Figure 6

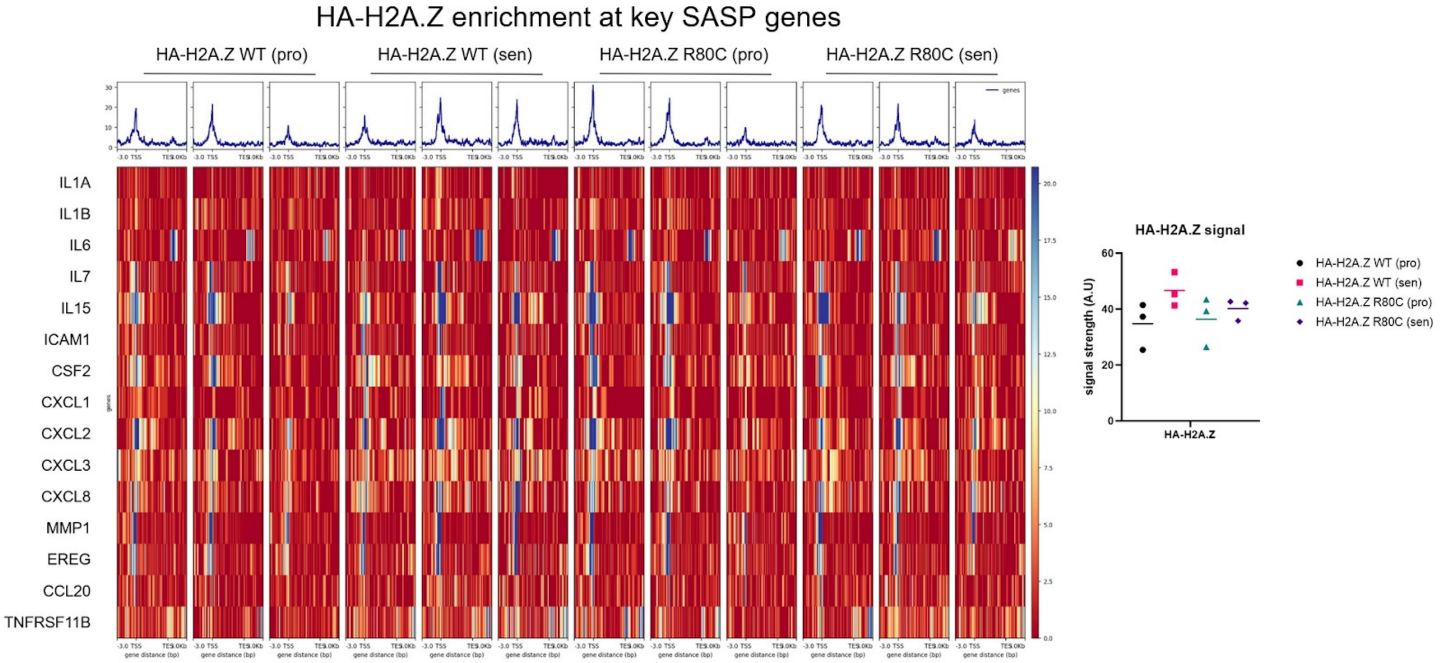
